# Brain-wide gaze-dependent activity during eyes-closed rest and sleep

**DOI:** 10.64898/2026.07.10.737751

**Authors:** Zachary Nudelman, Matthias Nau

## Abstract

Gaze behavior and brain activity are tightly coupled during perception and mental imagery. Whether this coupling extends to eyes-closed states remains largely unknown, primarily because measuring eye movements alongside brain activity is difficult when the eyes are closed. Here, we used a scalable MR-based eye-tracking approach to reconstruct gaze behavior from fMRI data during eyes-closed rest and sleep. We found widespread coupling between inferred gaze behavior and brain activity distributed through much of the cerebral cortex and cerebellum, including many regions typically associated with vision. These dynamics largely persisted across the two states. Additionally, accounting for gaze systematically altered functional connectivity estimates of large-scale brain organization. Together, these results underscore the importance of considering gaze as a hidden behavioral dimension of intrinsic fMRI signals, even in eyes-closed states. More broadly, our findings suggest that eye–brain dynamics may provide a window into eyes-closed states of consciousness.

## Introduction

Gaze behavior offers a powerful window into cognition and brain function. During wakefulness, eye movements and brain activity are tightly coupled to perception, imagination, and memory recall (1–3), suggesting that eye-brain dynamics can index cognitive states with or without visual input. Even in eyes-closed states such as sleep, links between eye movements and cognition have long been proposed, from the visual scanning hypothesis in rapid eye movement (REM) sleep (4–7) to neural evidence that REMs may reflect the content of dreams (8–13).

Yet, while pioneering efforts exist (8, 12–14), eye-brain dynamics in eyes-closed states remain poorly understood, in part because existing methods for measuring human gaze behavior in the MRI scanner are ill suited to this question: electrooculograms (EOG) require dedicated hardware and are prone to MRI-related artifacts, while camera-based eye tracking is typically unavailable in eyes-closed states entirely. Importantly, closing this gap could advance our understanding of eyes-closed states of consciousness such as dreaming (9, 15), and inform studies of disorders characterized by altered patterns of gaze and brain activity, including Parkinson’s disease and Alzheimer’s disease (16, 17).

Here, we address this gap using MR-based eye tracking, an approach that infers gaze behavior directly from the MR signal of the eyes (18–24). Unlike other eye-tracking approaches in MRI, our method can be applied retrospectively to existing datasets and does not require MR-compatible hardware or dedicated training data. This capability allowed us to analyze publicly available fMRI data collected while participants were falling asleep (25), and ask whether eye-brain dynamics persist even during eyes-closed states.

Using this approach in both eyes-closed rest and sleep, we found widespread gaze-dependent brain activity extending far beyond classical oculomotor regions to much of the cerebral cortex and cerebellum. This coupling was largely preserved across both states and prominently involved many regions typically associated with vision. Moreover, we show that accounting for inferred gaze systematically alters functional-connectivity estimates, a widely used measure in the study of large-scale brain organization. Together, these findings demonstrate the influence of gaze behavior on intrinsic brain activity in the absence of visual stimulation, suggesting that eye-brain dynamics indeed offer a window into eyes-closed cognition. Our MR-based eye-tracking approach further provides a scalable framework for probing eye–brain dynamics across states and populations that otherwise lack a continuous behavioral read-out, and pave the way for future studies of eyes-closed states of consciousness more broadly, including sleep, anesthesia, or coma.

## Results

In the following, we present our main findings in six steps. First, we introduce the fMRI dataset and our MR-based eye-tracking approach in detail, describing how time-resolved gaze predictors were inferred from eye-voxel patterns. Second, we relate these predictors to voxel-wise brain activity and show that gaze-dependent activity is widespread even in the absence of visual input, extending well beyond classical oculomotor and visual regions. Third, we compare eyes-closed rest and sleep to show that eye-brain dynamics largely persist across these states. Fourth, we show that accounting for inferred gaze dynamics systematically alters brain-wide functional connectivity. Fifth, we demonstrate that our results are not explained by head motion and align well with those obtained through EOG. Finally, we place these findings in the context of prior work on sleep and dreaming, as well as the use of resting-state activity in basic and clinical research.

### 1) Eyes-closed MR-based eye tracking reveals gaze-dependent brain activity

In this study, we analyzed fMRI data acquired during eyes-closed rest (hereafter referred to as Wake) and Sleep (25), including concurrently recorded fMRI, electroencephalography (EEG), and EOG data (Participants: n=33; fMRI, EEG, and EOG n=31 after exclusions; Figure 1A). Participants were instructed to try to fall asleep inside the scanner, and sleep staging was performed on the EEG data by a Registered Polysomnographic Technologist. Functional MRI data were preprocessed using fMRIprep with standard settings (*version 25.0.0*, (26), see methods).

**Figure 1:**
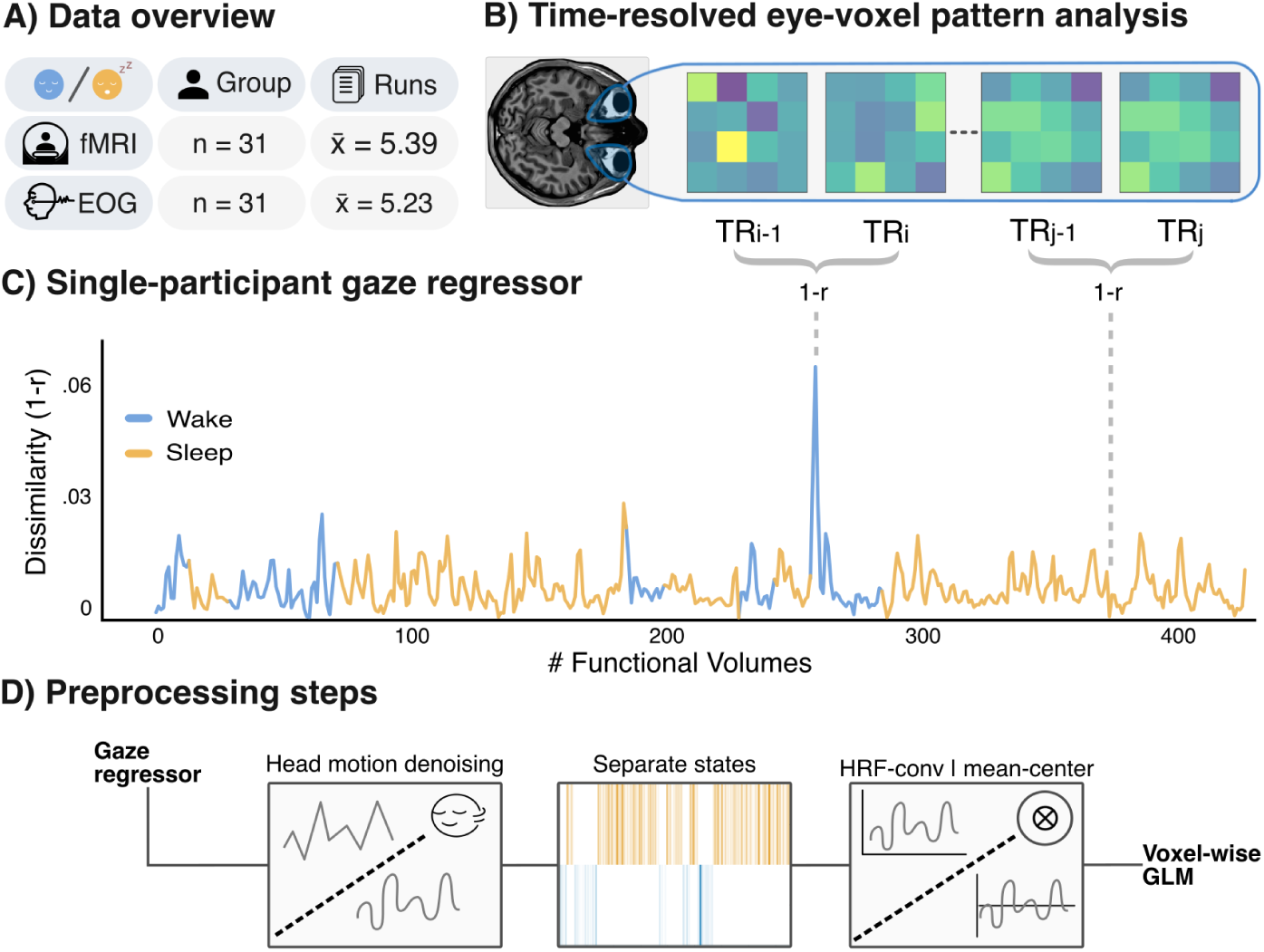
Inferring gaze from eye-voxel patterns during eyes-closed rest and sleep. (A) Dataset overview. Concurrent fMRI and EEG were recorded during 15-minute scanning runs in which participants (n=31) attempted to fall asleep. (B) Analysis logic. Gaze dynamics are inferred by comparing eye-voxel patterns across consecutive functional volumes using Pearson correlation. Greater dissimilarity reflects larger displacement of the eyes (1-Pearson’s r). (C) Single-participant example. Gaze predictor for an example participant (Participant 17 Run 3) separated into Wake (blue) and Sleep (orange). (D) Linking gaze to brain activity. The gaze predictor from each run was denoised with respect to head motion parameters, separated by state, convolved with the hemodynamic response function (HRF), and mean-centered. This procedure yielded a final gaze predictor which was then related to brain activity using a voxel-wise general linear model (GLM) analysis.

Building on previous work showing that eye-voxel patterns carry information about gaze position and movement (18, 19, 21, 22, 24), we developed a time-varying eye-voxel pattern analysis (MReye-Move) that reconstructs gaze behavior directly from the MR signal of the eyes. Unlike other eye-tracking approaches in MRI, this method allows for retrospective analysis of existing data without requiring MR-compatible hardware or training data. In brief, gaze behavior is inferred by comparing eye-voxel patterns across successive functional volumes (Figure 1B; for details, see methods). If the eyes remain stable, pattern similarity between volumes is expected to be high; if the eyes move, similarity is expected to decrease (Figure 1B–C). Thus, changes in eye-voxel patterns provide an index of gaze dynamics, quantified as 1-Pearson correlation between successive volumes, as shown in previous work ((24), Figure 1C).

Using this approach, we created a gaze predictor for each participant and scanning run, orthogonalized it with respect to head motion, split it into Wake vs. Sleep using EEG sleep staging, and then convolved the resulting regressor with the hemodynamic response function (HRF, (27), Figure 1D). This approach allowed us to examine the influence of gaze behavior on brain activity even though the eyes were closed, and thus in states where camera-based eye tracking is unavailable.

To determine the relationship between gaze behavior and brain activity, we fit a voxel-wise general linear model (GLM) to each brain voxel’s time series, with the HRF-convolved gaze predictor as the main regressor of interest (27). The GLM further included 16 nuisance regressors modeling head motion and their derivatives, framewise displacement, and global signal derived from cerebrospinal fluid, white matter, and the whole brain. These nuisance regressors were obtained during preprocessing with fMRIprep (26). In addition, constant terms were included to model the mean signal of each run. The resulting gaze-dependent brain-activity maps were averaged across runs, yielding one map per participant and state, which then entered the group-level analysis.

### 2) Wide-spread gaze-dependent brain activity during eyes-closed states

Examining the resulting gaze-dependent brain activity maps at the group level revealed that our gaze predictor covaried widely with brain activity in both states (one-sample t-test against zero, mixed-effects FLAME 1+2, FDR-corrected at *q* = 0.05, Figure 2A). Both positive and negative associations spanned across the brain, encompassing much of the cerebral cortex and cerebellum. For example, we found a pronounced positive relationship between inferred gaze behavior and brain activity in the occipital lobe as well as the oculomotor vermis, regions typically associated with visual processing and visuospatial cognition (28–30). In contrast, frontal and medial parietal cortices showed a negative relationship with inferred gaze behavior, likely overlapping with the default mode network (Figure S1), exhibiting lower activity when inferred eye-movement estimates were stronger.

To characterize the distribution of these effects across the visual system, we next performed a posthoc region-of-interest (ROI) analysis using atlas-defined retinotopic areas from the Benson atlas (28). Gaze-dependent activity was broadly distributed across these visual regions (Figure 2B), with most ROIs surviving FDR correction across regions and states (Table S1). Together, these results show that gaze-dependent brain activity is strikingly widespread during both eyes-closed rest and sleep, including many regions typically associated with vision. Critically, because the eyes were closed and participants were sleeping in key analyses, these effects occurred in the absence of external visual stimulation.

**Figure 2:**
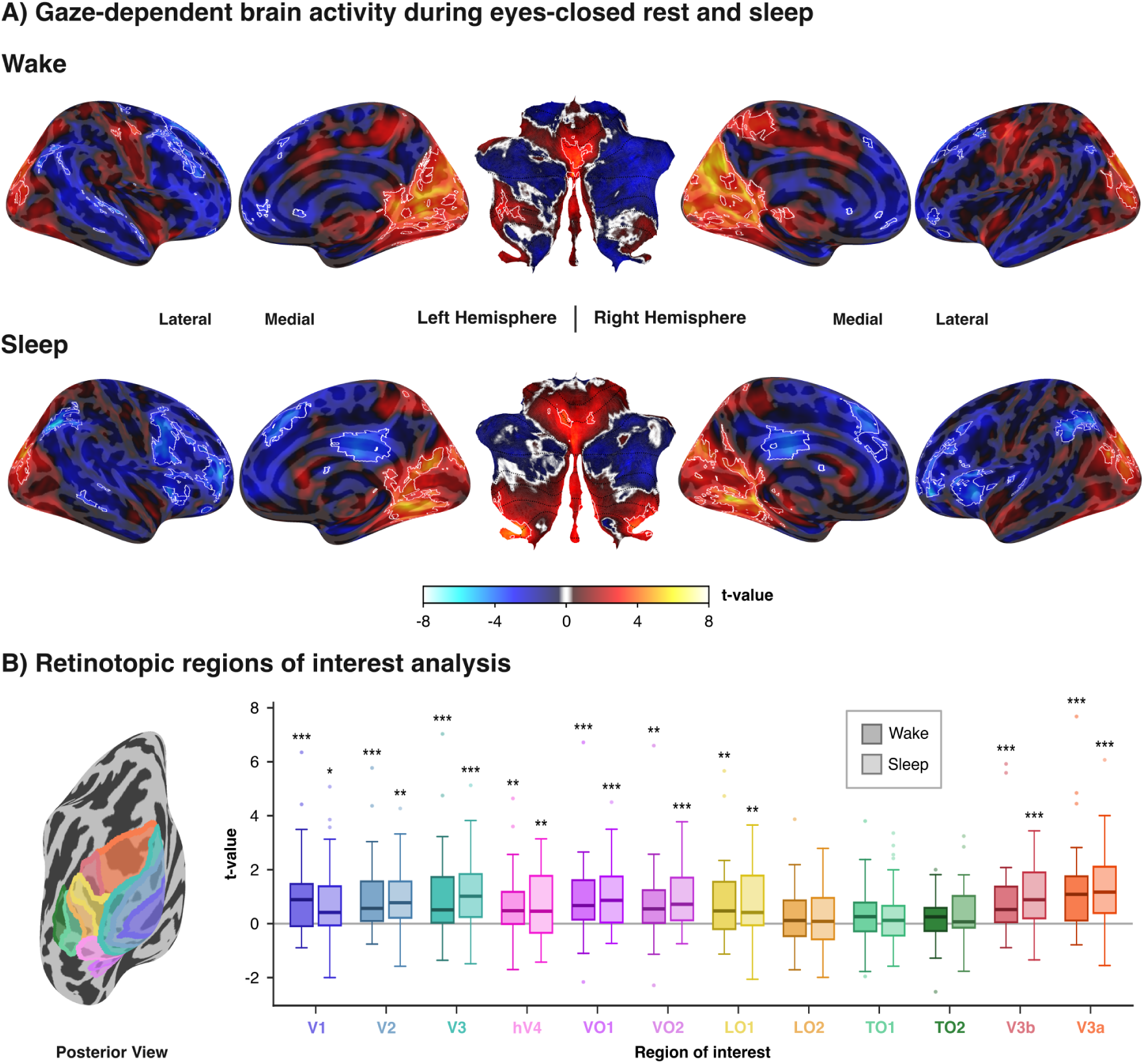
Widespread gaze-dependent brain activity during eyes-closed rest and sleep. A) Voxel-wise general linear model results estimated for the eye voxel-based gaze predictor in Wake (top) and Sleep (bottom). Maps display group-level results of one-sample t-tests performed against zero using FSL’s FLAMEO mixed-effects model (FLAME 1+2), with cerebral results projected on FreeSurfer’s FSaverage surface and cerebellar results mapped onto SUITPy’s flatmap surface. No t-thresholding was applied in order to show the global topographic organization of effects, with white contours indicating regions surviving False Discovery Rate (31) correction (*q* < 0.05, cluster extent > 20 voxels). B) Post-hoc regions of interest (ROI) analysis for atlas-defined retinotopic regions displayed on a posterior FSAverage surface (28). Shown are whisker box plots (central line: median; box: 25th and 75th percentile; whiskers: all data not considered outliers; outliers: data points outside 1.5×interquartile range), with color indicating ROI and statistics indicating results of sign-flipping permutation-based (10,000 permutations) two-tailed one-sample tests of subject-level t-scores against zero after false-discovery-rate correction (*q* = 0.05) across ROIs: * *p* < 0.05, ** *p* < 0.01, *** *p* < 0.001. See Table S1 for precise p-values. Eyes-closed gaze-dependent brain activity was distributed across regions involved in vision.

### 3) Gaze-dependent brain activity overlaps across rest and sleep

Having established gaze-dependent brain activity in both eyes-closed states, we next asked how these effects overlapped spatially, to determine whether similar brain structures were engaged in both states. We addressed this question by comparing the unthresholded group-level statistical maps across states using a searchlight-based pattern-similarity analysis (32) akin to earlier reports (24). This analysis quantified the multi-vertex similarity in gaze-dependent activity patterns across states, independent of whether the underlying effects were positive or negative. Because surface-based analyses better respect the topology of brain structure, we conducted these analyses using surface projections of our data (33).

Gaze-dependent brain-activity maps showed substantial spatial overlap across states (median vertex correlation: cerebrum *r* = 0.44, cerebellum *r* = 0.54; Figure 3A). Nevertheless, vertex-wise analyses also revealed localized state-dependent differences, most prominently in anterior parietal and superior temporal cortices, prefrontal cortex, and the occipital lobe (Figure 3A). However, none of the atlas-defined visual regions showed a significant difference between states, suggesting that the vertex-wise differences did not align with anatomically defined visual areas (28). These results indicate that gaze-dependent brain activity largely overlaps across eyes-closed rest and sleep, while revealing localized differences as key future targets for understanding state-dependent modulation (see Figure S1 for large-scale network-level ROI results).

Note that all previous analyses treated sleep as a single state, combining non-rapid eye movement (NREM) states NREM1 and NREM2. This approach increased the amount of available data and ensured that the number of samples as well as the mean amplitude and standard deviation of the gaze predictors were matched between Wake and Sleep (one-sample t-tests on pseudo-z scores from within-participant state-label permutation tests, all *p* > 0.05, *n* = 31; Figure 3B). However, NREM1 and NREM2 differ in the depth of sleep and potentially in their associated eye-brain dynamics. We therefore repeated our analyses separately for NREM1 and NREM2 in participants with sufficient data per state (n = 24). Gaze-dependent effects were strongest in NREM2 (Figures S2, S3), which further appears to drive the differences between Wake and Sleep observed earlier (Figure 3). These state-specific analyses suggest that eye-brain dynamics may in fact change as sleep deepens.

### 4) Eyes-closed gaze behavior shapes estimates of large-scale brain organization

Given the distributed nature of gaze-dependent brain activity in our data (Figure 2), we next asked whether accounting for gaze influences measures of large-scale functional brain organization. In the absence of an explicit task or stimulus, such organization is commonly characterized using functional connectivity (FC), which captures covariation in activity between brain regions. To test whether eyes-closed gaze behavior shapes FC estimates, we computed pairwise Pearson correlations between z-scored time series from 500 Schaefer atlas parcels (34), either before or after removing gaze-related effects. Both raw and HRF-convolved gaze predictors were included to remove instantaneous and hemodynamic gaze-related signals, respectively. In all cases, we additionally included our 13 nuisance regressors modeling head motion as before, as well as global signals derived from cerebrospinal fluid and white matter. In this analysis, all states were modeled together to maximize the amount of available data (35). After estimating parcel-wise FC, the resulting estimates were grouped by the Yeo-7 networks for visualization and overview (Figure 4). These networks comprise regions with similar connectivity profiles during rest (36).

**Figure 3:**
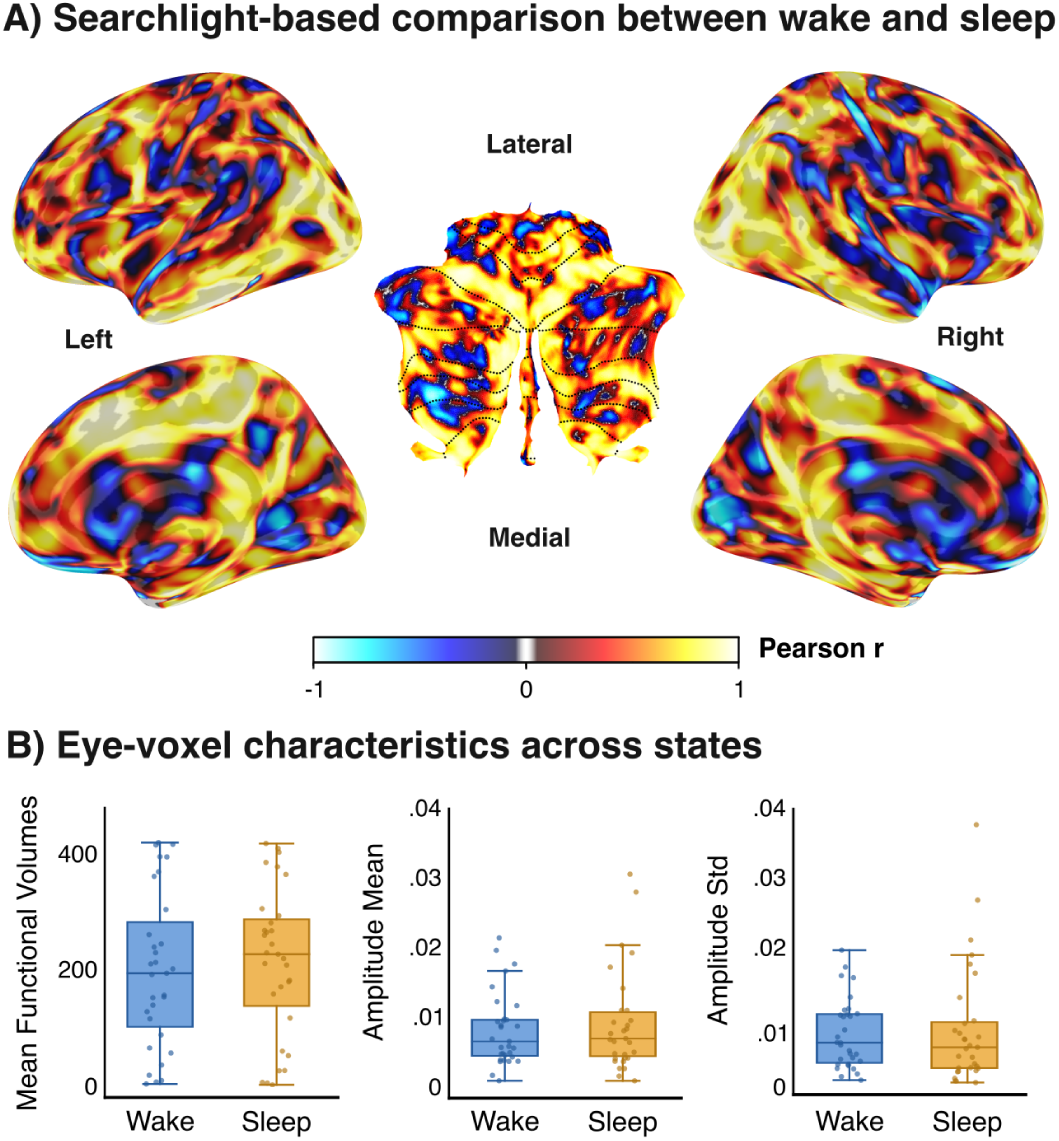
Across-state comparison between Wake vs. Sleep. (A) Searchlight-based local similarity reveals widespread overlap between Wake and Sleep. Statistical maps show the local similarity between the group-level t-statistic results for Wake and Sleep shown in Figure 2. The maps were compared using Pearson correlation (r) within local geodesic searchlights of 23*mm* radius for the cerebral cortex and 11*mm* for the cerebellum. Warm colors indicate spatial consistency across states, whereas cool colors indicate localized differences. (B) Comparison of gaze metrics across states. Left panel: across-run mean number of functional volumes in each state. Center and right panels: group-level mean and standard deviation of gaze predictor amplitude, respectively. Dots represent individual participants. No significant differences were found between Wake and Sleep for any of these metrics (one-sample t-tests on pseudo-z scores from within-participant state-flipping permutation tests, all *p* > 0.05).

We found that gaze behavior influenced FC estimates across all networks (see Figure 4), with around 11% of all parcel-to-parcel edges surviving correction (sign-flipping permutation test, 10,000 permutations, FDR correction at *q* < 0.05). Of the significant edges, 83% showed reduced FC after accounting for gaze, mostly within visual and somatomotor networks, and between the dorsal attention network and all others. Conversely, 17% of significant edges showed increased FC after accounting for gaze, particularly between parcels with an opposing sign of gaze-dependent activity (Figure 2). Notably, this included edges between visual and default mode network parcels known to be anticorrelated during rest (37). These results show that eyes-closed gaze behavior systematically shapes both network-level brain activity (Figure S1) and covariation among brain regions (Figure 4).

**Figure 4:**
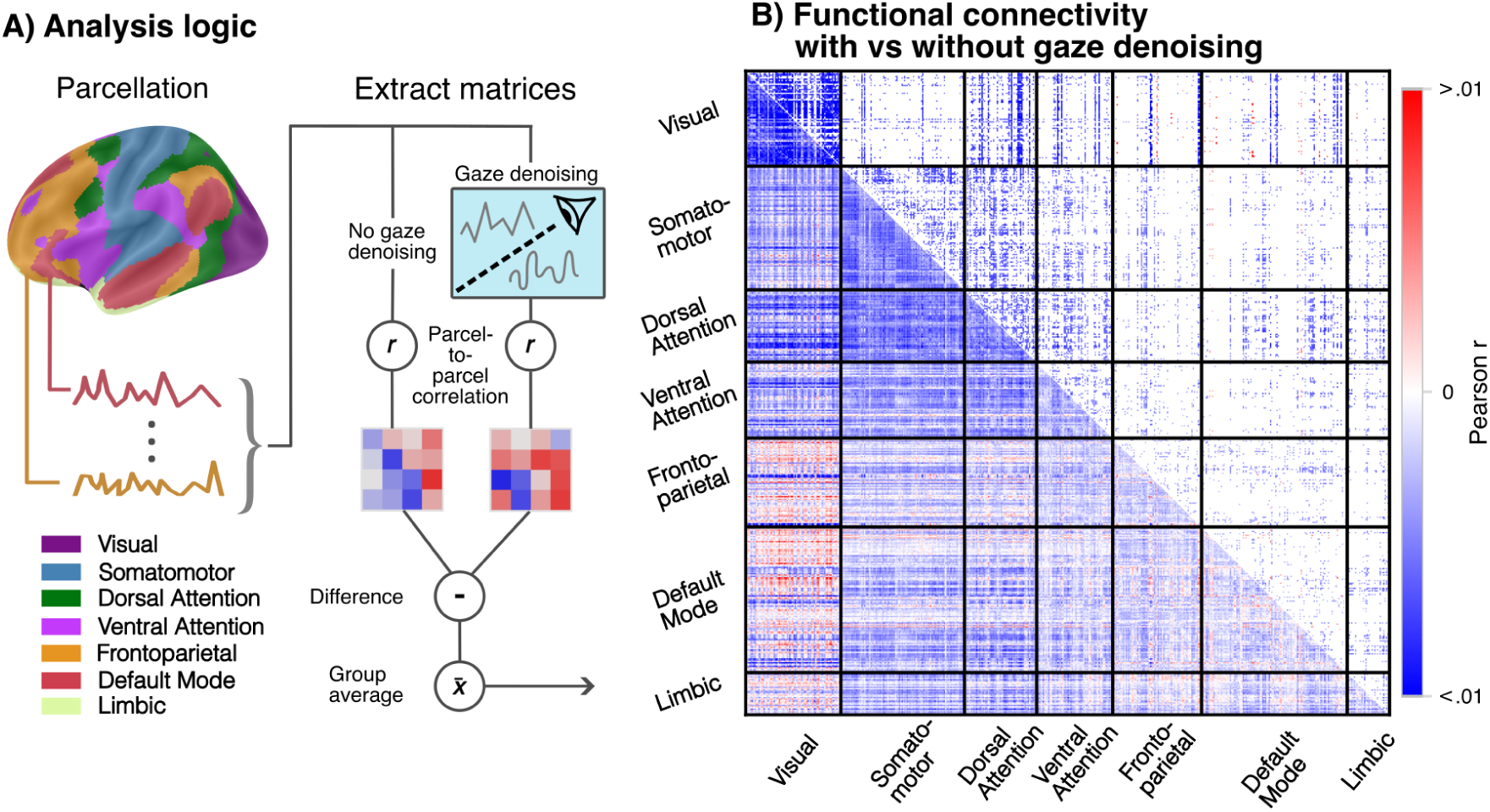
Eyes-closed gaze behavior shapes estimates of large-scale brain organization. (A) Analysis logic. BOLD time series were parcellated using the Schaefer 500-parcel atlas (34). Parcel-to-parcel Pearson correlation matrices were computed with and without removing variance explained by the raw and HRF-convolved gaze predictor. (B) Group-level edge-wise differences in functional connectivity. The lower triangle displays the unthresholded group-level mean differences in Pearson correlations between with vs. without gaze denoising; the upper triangle displays edges surviving False Discovery Rate correction at *q* = 0.05, two-sided sign-flipping permutation test, 10,000 permutations. Red values indicate higher functional connectivity after gaze denoising of each voxel’s time series, with blue indicating the opposite. Results were grouped by the Yeo 7-networks for overview (36), with each network comprising many parcels.

### 5) MR-based eye tracking captures gaze dynamics

To rule out alternative explanations, we tested whether our MR-based gaze predictor revealed meaningful brain dynamics related to gaze rather than nonspecific image fluctuations. First, we asked whether the effects could be driven by head motion. While the instantaneous effects of head motion were modeled in all analyses (i.e., head-motion parameters, their derivatives, and framewise displacement), we additionally implemented three distinct denoising strategies: 1) No additional head-motion denoising, 2) Modeling the hemodynamic effects associated with head motion in the voxel-wise GLMs, 3) Orthogonalizing the gaze predictor with respect to head motion (Strategy 3 reflects our main analyses presented earlier). Importantly, our main results were highly robust across all denoising strategies (Figure S4), suggesting that any potential effect of head motion on gaze-dependent brain activity was minimal (see methods for details).

Second, we asked how our eye voxel-based approach compared to those obtained through concurrently recorded EOG. To construct an EOG-derived gaze predictor, we computed the magnitude of the framewise difference in mean EOG signal across functional volumes, similar in logic to our eye-voxel predictor (see methods). Despite different noise sources and measurement principles of eye-voxel signals and EOG (38), the two regressors were correlated (*r* = 0.416, average across participants; one-sample t-test: *p* < 0.001; Figure S5C), indicating substantial shared variability. Moreover, fitting the same GLM with the EOG gaze predictor showed a spatial pattern highly similar to the eye-voxel-based results (Figure S5A). Finally, quantifying the overlap in eye voxel-based vs. EOG-based gaze-dependent brain activity confirmed their high correspondence (combined Wake and Sleep median vertex correlation: cerebrum *r* = 0.67; cerebellum *r* = 0.72; Figure S5B). Together, these analyses provide strong evidence that our gaze predictor indeed captured meaningful gaze dynamics.

## Discussion

The present study shows that brain activity covaries widely with gaze behavior even in the absence of visual input. These eye–brain dynamics encompassed much of the retinotopically-defined visual system and were largely preserved across eyes-closed rest and sleep. Moreover, we report evidence that eyes-closed gaze behavior affects brain-wide functional connectivity estimates, which form the basis for many studies of the human brain at rest. Our findings were robust across denoising strategies, including multiple controls for head motion, and closely aligned with results obtained from concurrently recorded EOG. Together, these findings establish eye-brain dynamics as a window into eyes-closed states of cognition, and identify gaze behavior as an underappreciated contributor to intrinsic fMRI signals. More broadly, our work highlights the importance of measuring gaze behavior in fMRI studies, and establishes an MR-based eye-tracking method that can be used without training data or MR-compatible equipment, and even when the eyes are closed. Because the approach is broadly applicable and works in existing data, it may further enable new, large-scale studies of eyes-closed states of consciousness.

In the following, we discuss our findings in light of previous work on dreaming and eyes-closed cognition, before exploring how gaze behavior and MR-based eye tracking can make both resting state and task-based fMRI studies more “naturalistic”.

### Gaze behavior as a window into eyes-closed cognition?

The present work was motivated by the idea that eye-brain dynamics track cognitive processes even in the absence of visual input (for review, see (1–3, 9, 15)), and that these dynamics may extend to eyes-closed states such as mind wandering and dreaming. Indeed, during awake recall of previously seen stimuli, spontaneous eye movements have been shown to mimic the spatial layout of imagined or remembered stimuli, even while looking at a blank screen or sitting in complete darkness (24, 39–43).

Extending these ideas to sleep, eye movements during REM sleep have long been proposed to provide a window into dream imagery under the visual scanning hypothesis (4, 5). Some of the strongest evidence comes from patients with REM sleep behavioral disorder, who lack normal REM atonia and often enact their dreams: in these individuals, the direction of rapid eye movements is frequently aligned with the direction of dreamed actions (7). In rodents, parallel findings show that rapid eye movements during REM sleep signal the direction and amplitude of ongoing changes in virtual head direction (10). Our results are consistent with these earlier reports. Critically, since eye-brain dynamics reflect visual perception and mental imagery during wakefulness (1–3), and eye movements during wakefulness and sleep engage classical visual regions (Figure 2, (13, 14)), we speculate that these dynamics reflect subjective experiences arising during eyes-closed mind-wandering and sleep. Studying eye–brain dynamics across cognitive states may therefore help elucidate the general principles by which experiences are constructed across visual perception, imagery, and dreaming (44, 45).

An important alternative interpretation is that our results reflect oculomotor signals, such as efference copies, acting on brain-wide activity independent of dreaming. While this interpretation would itself be interesting, at least three points speak against it. First, when the eyes are open, efference-copy signals are thought to suppress visually evoked responses rather than driving gaze-dependent activity as observed here (46–48). Second, the localized state-dependent effects we observed in our voxel-wise analyses (e.g., group-level results were robust in the calcarine sulcus only during Wake, not Sleep, Figure 2), suggest that the relationship between gaze behavior and activity could be dynamic and unlikely reflects a purely motoric signal. Our separate analyses of NREM1 and NREM2 further support this idea (Figure S2). Finally, when the eyes are closed, guided eye movements do not evoke gaze-dependent brain activity in visual cortex (14, 49), while spontaneous eye movements do (Figure 2, (49)). Such inconsistencies and state-dependent effects are difficult to explain by oculomotor signals alone. While future work must further establish whether eye movements indeed reflect dream content, oculomotor signals, or co-varying factors such as arousal (50), our findings align with the broader view that gaze and brain activity jointly reflect eyes-closed cognition such as mind wandering and dreaming (4, 5, 9, 15).

### MR-based eye tracking for naturalistic studies of rest and sleep

A key methodological decision when designing task-free fMRI experiments is what the participant should do with their eyes: keep them closed, free viewing with eyes open, or eyes-open fixation. Intrinsic activation patterns and connectivity estimates vary across these conditions (51–56), possibly due to between-participant differences in gaze behavior when fixation is not enforced (Figure 2). In line with this idea, test-retest reliability appears to be highest during fixation and lowest when the eyes are closed (54). For this reason, many resting-state studies enforce fixation, which increases test-retest reliability and functional connectivity estimates of visual networks with other networks, possibly due to the absence of eye movements and visual motion (55).

However, we question whether such behaviorally constrained states can still be considered natural “rest”, and whether the fixation task itself may affect brain activity and other read-outs of cognition. While fixation may enhance experimental control and group-level statistics, it does so by suppressing spontaneous behavior that may itself be integral to participants’ cognition and brain activity that studies seek to understand. For example, eye movements have been linked to mental imagery, recall, and dreaming (1–3, 15), but enforcing fixation reduces vividness of such internal experiences (e.g., (57, 58)). Fixation during resting state is therefore likely to affect in-scanner cognition and the functional-connectivity patterns associated with it (59)). We argue that rather than suppressing or ignoring eye movements during resting state, gaze behavior should be measured and included in the analysis, as it contributes meaningful variance within and across brain regions both in eyes-open (23, 60, 61) and eyes-closed paradigms (Figures 2,4). More broadly, gaze should be treated not as a nuisance variable, but rather as a meaningful behavioral signal in its own right; one that can render both task-based and resting-state paradigms more “naturalistic” (45, 62, 63).

This argument naturally extends to other behaviorally unconstrained states, including sleep. Using our approach (Figure 1), gaze can be included as one of many physiological measurements whose shared dynamics may be crucial in understanding the functions of sleep (64). Although the effect of gaze on functional connectivity estimates was modest for the metric examined here (Figure 4), it was robust at the group level, and distributed throughout the brain, suggesting that eye–brain dynamics exert a systematic influence on such large-scale network organization estimated with fMRI. Indeed, when accounting for gaze, the well-established anti-correlation between sensorimotor regions and the default mode network weakened (Figure 4B, (37)). More broadly, our approach could complement studies of occipital responses to visual stimulation during sleep (65) as well as hemodynamic and metabolic changes across sleep stages (66, 67), by offering an additional behavioral dimension with which to interpret brain activity. This may be particularly useful given our exploratory evidence that eye–brain dynamics themselves seem to vary across sleep stages (Figure S2).

### Inferring gaze behavior from eye-voxel patterns

The most common method of measuring eyes-open gaze behavior is camera-based eye-tracking. For eyes-closed paradigms, some studies have turned to EOG-based eye tracking (49, 61) or video recording participants’ faces and limbs to determine onset and duration of rapid eye movements (12). While these approaches are valuable, they have important limitations, including the need for dedicated hardware and susceptibility to MRI artifacts in EOG, or the labor-intensive nature of manually labeling video. By contrast, our MR-based eye tracking approach requires no additional MR-compatible hardware and can be used post-hoc in existing data, yet it recapitulates the results obtained through these other methods (Figure 2, Figure S5, (12, 22, 49, 61)). MR-based eye tracking therefore presents the most scalable solution for examining eye-brain dynamics in eyes-open and eyes-closed states in fMRI.

Note that our method builds on previous approaches, which have already demonstrated that eye-voxel patterns contain information about the position and movement of the eye (18–22, 68–71). For example, gaze position has been successfully decoded from eye voxels during guided saccades, smooth pursuit, and visual search (21, 22, 24), while gaze position and, to a lesser degree, saccade parameters such as amplitude and duration have been linked to eye voxel patterns during movie viewing (24, 71). Moreover, related approaches linking the mean MRI signal of the eye to brain activity showed that gaze explained variance across many brain regions during eyes-open resting state (23, 60), even in blind individuals (72), which aligns well with our results obtained in eyes-closed rest and sleep (Figure 2, Figure 4). Notably, gaze decoding approaches have already yielded evidence that MR-based eye tracking may be feasible in eyes-closed states, as decoded eye-ball orientation matched participants’ self-reports of how their eyes moved (22).

Our results align with these earlier findings while extending them in important ways. First, to our knowledge, this study provides the first application of MR-based eye tracking showing gaze-dependent brain activity during eyes-closed rest and sleep, states that are inherently difficult to assess with conventional eye tracking. Second, unlike prior MR-based gaze-decoding approaches, our method does not require dedicated training data, making it scalable and readily applicable to existing fMRI datasets. Third, by demonstrating that eye–brain dynamics can be studied when the eyes are closed, our findings expand the scope of MR-based eye tracking beyond guided eye movements and wakeful viewing paradigms. More broadly, this approach may enable the study of eye–brain dynamics across eyes-closed states of cognition, including sleep, coma, anesthesia, or disorders of consciousness

### Limitations and open questions

One limitation of the present study is the absence of a direct “ground-truth” measure of gaze behavior. For example, despite instructions to fall asleep, and the presence of sleep-related EEG rhythms, occasional eye openings cannot be fully excluded, even during sleep. However, such events are expected to be sporadic and highly idiosyncratic, making them unlikely to account for the robust and spatially distributed group-level effects observed in the present study (Figure 2). Relatedly, apart from EOG recorded from a single electrode, no independent measure of eye movements was available. However, previous work has successfully validated comparable MR-based eye-tracking approaches against camera-based eye tracking in eyes-open states (19, 21, 22, 24) and against self-reports of guided eyes-closed eye movements (22). Moreover, our results aligned closely with those obtained using EOG (Figure S5) and with previous reports of REM-related, NREM-related, and eyes-open brain activity (12, 23, 49, 60). Together with our own EOG analyses (Figure S5), this close alignment with previous work supports the notion that the present gaze predictor indeed captured biologically meaningful oculomotor dynamics. Future work combining more extensive EOG recordings and instructed eyes-closed eye movements could help clarify the precise factors that drive eye-voxel pattern similarity differences across functional volumes (e.g., few large eye movements vs. many small ones).

Importantly, one factor that could drive changes in eye-voxel patterns is head motion. For that reason, we removed variance explained by head motion from the gaze predictor in all main analyses (Figure 1), and we conducted extensive control analyses to rule out the effects of head motion on our gaze-dependent brain activity estimates (Figure S4). Moreover, it is not obvious why head motion would selectively affect the regions showing gaze-dependent activity in our data (Figure 2), such as the occipital lobe. That said, we note that head motion, like gaze, could reflect ongoing cognition in a meaningful way (73), and that head motion and gaze may genuinely covary during natural behavior. Indeed, eyes-open gaze shifts and head movements often align (74), making it difficult to disentangle their distinct associations with cognition.

An interesting open question that remains is whether eyes-closed eye-brain dynamics reflect the content of dreams or other aspects of cognition during sleep. To address this question, future work could compare dreaming versus non-dreaming periods (75), or directly relate fluctuations in eye-voxel patterns and associated brain activity to dream reports (76), dream-content decoding (77, 78), or markers of memory reactivation (79–81). Ideally, such studies would involve longer sleep sessions that permit a broader sampling of sleep stages including REM (8), which is particularly interesting in light of the evidence for differences between NREM1 and NREM2 (Figure S3). A particularly fascinating avenue would be to study lucid dreamers (82), in whom eye movements have already been used to gain insights into the neural correlates of dreams (83–85).

More broadly, comparing gaze-related neural mechanisms across many eyes-open and eyes-closed states may help clarify long-standing debates regarding the conceptual boundaries between dreaming, perception, and mental imagery (15, 45, 85–87). To address such broad questions, MR-based eye tracking may prove valuable, as it can be applied to cognitive states and populations that are otherwise difficult to study. Particularly promising applications include disorders of consciousness, coma, anesthesia, and other minimally responsive states (17, 88–90).

## Conclusion

In conclusion, by linking eye-voxel signals to brain activity measured with fMRI, we show that gaze and brain activity covary widely even when the eyes are closed. The spatial distribution of gaze-dependent brain activity was largely preserved across eyes-closed rest and sleep, and accounting for these effects systematically altered functional connectivity estimates throughout the brain. These findings suggest that eye–brain dynamics provide a window into eyes-closed states of cognition, establish MR-based eye tracking as a scalable approach for studying these states, and high-light the importance of considering gaze when interpreting fMRI data even in the absence of visual input.

## Data availability

The fMRI data used in this work were shared by the original authors (25) and can be downloaded from openneuro.org: https://openneuro.org/datasets/ds003768/versions/1.0.13.

## Code availability

Python code underlying our key analyses is available in the following GitHub repository: https://github.com/BCN-lab/SleepMReye. MReyeMove, our MR-based eye-tracking pipeline, is currently being prepared for release in a user-friendly format through the OpenMReye GitHub organization: https://github.com/deepmreye.

## Acknowledgements

We thank Wietske van der Zwaag, Freek van Ede, and Chris Baker for helpful feedback on our manuscript draft, as well as Serge Dumoulin and Tomas Knapen for helpful discussions. We also thank Gu and colleagues for making their data publicly available.

## Author contributions

MN conceptualized the present study, guided the research design and interpretation of findings, and provided overall supervision. ZN performed the data analysis, generated the figures, and wrote substantial portions of the manuscript. MN wrote and revised substantial portions of the manuscript. Both authors reviewed and edited the manuscript, approved the final version, and agree to be accountable for the work.

## Ethics declaration

The authors declare no competing interests.

## Methods

### 1) Participants

Publicly available functional magnetic resonance imaging (fMRI) data of N=33 participants (age: 22.1±3.2 years, female/male: 16/17) were downloaded from openneuro.org (https://openneuro.org/datasets/ds003768). These data were released as part of an earlier report (25). The study was approved by the Institutional Review Board at the Pennsylvania State University (Protocol numbers: STUDY00005969 and STUDY00015305) and participants gave written informed consent prior to scanning. Two participants were excluded because of a lack of sleep data (sub-02, sub-21). Across all participants, a total of 10 scanning runs were excluded due to excessive head motion (exceeding a mean framewise displacement of 0.5 mm), in addition to another 12 runs that were excluded because sleep staging was incomplete. Data of 31 participants and a total of 167 scanning runs entered analysis.

### 2) Experimental procedure

The experiment consisted of the following phases reported in chronological order: a 2-min EEG quality-control scan; a 5-min T1-weighted structural scan; a 10-min resting-state scan; a 15-min visual-motor task scan; another 10-min resting state scan; followed by several 15-min sleeping scans (25). Note that only the sleep scans were considered in the present work in which participants were instructed to try to fall asleep. During these runs, participants indicated their level of wakefulness by clicking a button approximately every second, stopping when transitioning to sleep.

### 3) fMRI acquisition and preprocessing

Functional MRI data were acquired on a Siemens 3T Prisma Fit scanner using a 20-channel receiver head coil at Pennsylvania State University. T1-weighted structural scans were acquired using a MPRAGE sequence (TR = 2300 ms, TE = 2.28 ms, TI = 900 ms, flip angle = 8°, FOV = 256 mm, matrix size = 256 x 256 x 192, voxel size = 1 x 1 x 1 mm, acceleration factor = 2, phase-encoding direction along the A-P axis). BOLD functional images were collected with an echo-planar imaging (EPI) sequence (TR = 2100 ms, TE = 25 ms, flip angle = 90°, 35 interleaved axial slices, slice thickness = 4 mm, FOV = 240 mm, in-plane resolution = 3 x 3 mm).

Data were preprocessed using *fMRIPrep* 25.0.0 (26?). Anatomical images were corrected for intensity non-uniformity using N4BiasFieldCorrection distributed with ANTs 2.5.4 (91). Cerebrospinal fluid (CSF), white-matter (WM) and gray-matter (GM) were segmented on structural scans using fast (92). The scans were normalized to the MNI152NLin2009cAsym space by nonlinear registration using antsRegistration. Slice-timing correction was applied, with slices interpolated to the middle of the TR (start time = 1.01 s). Head motion parameters for all runs of each participant were estimated with mcflirt (93) using a reference image constructed by *fMRIPrep* to yield the six corresponding rotation and translation parameters. Temporal derivatives of these were also computed (94). Framewise displacement (FD) was calculated as the sum of the absolute magnitude of the first derivatives of the six head motion parameters (95). Functional data were co-registered to the processed anatomical reference image with mri_coreg (FreeSurfer 7.3.2) and flirt (96) using a boundary-based registration (97) with 6 degrees of freedom. They were then head motion corrected with the estimated head motion parameters.

### 4) Electroencephalogram (EEG)

EEG data were recorded using a BrainAmp MR-compatible 32-channel EEG system with a sampling rate of 5000 Hz and a passband of 0-250 Hz. An EOG electrode with an impedance below 20 kΩ was placed under the left eye and referenced relative to the midline fronto-central EEG electrode (FCz), with an impedance below 10 kΩ for the duration of the experiment.

### 5) Sleep Staging

Sleep staging was performed by a Registered Polysomnographic Technologist following the American Academy of Sleep Medicine (AASM) recommendations on acquired EEG data from the F3, F4, C3, C4, O1, and O2 channels (98). These data were preprocessed to remove MR gradient and ballistocardiogram artifacts (99), re-referenced to the contralateral mastoid (with ipselateral or averaged mastoids used as a fallback for artifact affected channels), and band pass filtered between 0.3-35 Hz. Sleep staging into Wake or Sleep (NREM1, NREM2, NREM3) was then performed on 30s epochs, with each epoch corresponding to 15 functional volumes. Further details of the EEG acquisition and staging procedure can be found in (25). Note that Sleep was defined in the present study as a combined state that included NREM1 and NREM2. Functional volumes staged as NREM3 were ignored in the analyses.

### 6) Gaze-predictor design

A gaze predictor was constructed using a time-resolved multi-voxel pattern analysis approach similar to that of Nau and colleagues ((24), Figure 1). First, eye voxels were extracted using an established and automated pipeline implemented in DeepMReye v0.3 (22). This pipeline co-registered each participant’s functional data of each run non-linearly to the DeepMReye template space in which the eyes were delineated manually. It then further co-registered a facial bounding box and finally the eyes specifically to those in the template, yielding the multi-voxel pattern of the eyes for each functional volume.

To infer gaze dynamics, our analysis relied on the following logic. If the eyes remained stable across two consecutive volumes, the corresponding eye-voxel patterns should be highly similar. Conversely, if the eyes moved between volumes, this pattern similarity should decrease. Following this logic, a gaze predictor was constructed by computing the dissimilarity between subsequent volumes, defined as one minus the Pearson correlation.

To remove any potential effect of head motion on the gaze predictor, we residualized it with respect to the six rigid-body motion parameters, their derivatives, and framewise displacement using Nilearn’s *signal.clean* function (100). This procedure yielded a signal linearly uncorrelated with head motion. Occasional negative deflections in the regressor after residualization were clamped to 0. See Section “Head motion control analyses” for alternative denoising strategies, all yielding highly similar results (Figure S4).

Because each dissimilarity value reflected the change between two adjacent volumes, it was assigned to the temporal midpoint between them. To align the gaze predictor with the fMRI volume acquisition times, we shifted it onto the TR grid using two-point moving-average interpolation. This procedure yielded a regressor that is two volumes shorter than the original voxel time series; accordingly, the first and last volumes of each scan were removed to maintain temporal alignment.

### 7) Linking gaze predictors to brain activity

We related the gaze predictor to brain activity using a mass-univariate general linear model (GLM). For each scanning run, the gaze predictor was split into two separate regressors corresponding to the conscious states Wake and Sleep. Each state-specific gaze predictor was then convolved with the hemodynamic response function (HRF) as implemented in Statistical Parametric Mapping (SPM) before mean centering.

Beyond the regressors of interest, the GLM included the six rigid-body motion parameters as well as their derivatives, framewise displacement, cerebrospinal fluid, white matter, and the global signal as nuisance regressors. Moreover, slow signal drifts were modeled using a cosine drift basis with a high-pass cutoff of 0.01 Hz, together with an intercept. Because constructing the gaze predictor removed the first and last time points, both the confound regressors and the functional images were trimmed as well to maintain temporal alignment.

The first-level GLM was fit using Nilearn’s *FirstLevelModel* (100) with a spatial smoothing kernel of 6 mm FWHM and an autoregressive noise model (AR(1)). For each participant, fixed effects were computed across runs using Nilearn’s *compute_fixed_effects* function to obtain contrast (COPE) and variance (VARCOPE) maps. These maps were then passed to a mixed effects Bayesian FLAME 1+2 model implemented in FMRIB Software Library (FSL) (101) using Nipype (?) to estimate the group-level effects.

For visualization, group-level maps were projected onto FreeSurfer’s FSAverage cerebral surface (102) using Nilearn. The cerebellum was projected onto a surface-based flatmap with SUITPy (using FSL space) applied directly to the MNI-aligned volumetric maps (103). Rather than thresholding these group-level maps, we visualized the unthresholded maps, with contours depicting significant clusters surviving FDR correction (*q* < 0.05, > 20 voxel extent).

Note that several runs had incomplete cerebellar coverage in the fMRI acquisition. Cerebellar analyses were therefore restricted to runs in which at least 90% of cerebellar voxels were available, as defined using the FSL Cerebellum-MNIfnirt probability map (thresholded at 25%, 2 mm resolution). This criterion retained 138 of 167 runs from 29 of 31 participants. The cerebellar probability map is distributed with FSL and was derived from the probabilistic atlas of Diedrichsen et al. (104).

To understand potential differences between the cognitive states in more detail, we additionally repeated the same GLM pipeline using the three states Wake, NREM1, and NREM2. However, the number of functional volumes per state was unbalanced across participants (per-participant mean TRs per run; median [IQR] across participants: W = 197 [104.33 - 285.57]; Sleep (NREM1 + NREM2) = 230 [140.43 - 290.42]; NREM1 = 128.00 [71.46 - 194.08]; NREM2 = 49.67 [6.58 - 122.33]). Therefore, we limited the main analysis to comparing Wake and Sleep as two broader states for which the amount of data was approximately balanced (Figure 3C).

### 8) Searchlight-based correlation analysis

The group-level gaze-dependent brain activity maps obtained for each cognitive state were compared in terms of the spatial distribution of effects using a searchlight-based correlation analysis (32). These analyses were performed on the cortical and cerebellar surfaces in order to better respect the brain’s folded anatomical structure (33). The geodesic neighborhood of each vertex on the cortical surface was computed with Dijkstra’s shortest path algorithm. The neighborhood radius was selected to maximize the median Jaccard overlap, across 10,000 randomly placed vertex centers, between surface neighborhoods and co-centered 3-voxel-radius volumetric search-lights projected onto the cortical surface, corresponding to a typical searchlight size in volumetric searchlight-based correlation analyses. This yielded an optimal surface radius of 23 mm for the cortex. Group cerebellar flat maps were also compared using the same procedure. However, a 2-voxel volumetric searchlight reference radius was used to find an optimal neighborhood size, reflecting the tighter folding of the cerebellum relative to the cerebrum (103). This procedure yielded a radius of 11 mm for the cerebellum. The value of each vertex was then calculated as the Pearson correlation across vertices within the geodesic neighborhoods between the two surface maps.

### 9) Head-motion-control analysis

Because we residualized our gaze predictor with respect to head motion in our main analyses (Figure 2), it was by definition uncorrelated with head motion. However, because the unprocessed gaze predictor was moderately correlated with framewise displacement before this residualization procedure (group-level mean correlation *r* = 0.281, 95% CI [0.224, 0.335], n=31), we conducted additional control analyses to understand the effect of head motion on our results in detail.

In addition to our main analyses that used a denoised gaze predictor, we repeated both the first-level and the second-level analysis using an “uncleaned” gaze predictor, which included the variance explained by head motion. The resulting gaze-dependent brain activity maps showed a highly similar pattern of results (unprocessed vs preprocessed combined state searchlight-based correlation of median vertex: cerebrum *r* = 0.90, cerebellum *r* = 0.92; see Figure S4). When comparing Wake vs. Sleep in terms of framewise displacement amplitude, we found a difference between states (Wake - Sleep = 0.05 mm, one sample t-tests on pseudo z-scores from within participant state-flipping permutation test, *t*(30) = 3.26, *p* = 0.0027, n = 31). However, this difference did not translate to gaze predictor amplitude differences even in our unprocessed gaze predictor (see Section 3, Figure 3C), nor did it differentially affect gaze-dependent brain activity estimates within each state (median vertex correlation between analyses for Wake: cerebrum *r* = 0.92, cerebellum *r* = 0.92; and Sleep: cerebrum *r* = 0.91, cerebellum *r* = 0.93).

Finally, we repeated the full pipeline again using the “uncleaned” regressor, this time modeling the hemodynamic effects of head motion by including an HRF-convolved framewise-displacement regressor as a covariate. Note that this regressor was added in addition to the six rigid-body motion parameters, their derivatives, instantaneous framewise displacement, cerebrospinal fluid, white matter, and the global signal that were included in all analyses. Again, even when removing the hemodynamic components of head motion, the analysis yielded highly similar gaze-dependent brain activity maps to our main analysis (median vertex correlation in Wake: cerebrum *r* = 0.96, cerebellum *r* = 0.92; in Sleep: cerebrum *r* = 0.97, cerebellum *r* = 0.93). Together, these control analyses show that our main results were extremely robust across head-motion denoising strategies, indicating that our results are unlikely to be explained by head motion.

### 10) Electrooculography (EOG)

To examine an alternative measure of gaze behavior, we constructed an EOG-based gaze predictor similar in logic to our eye-voxel based one. The underlying EOG data were recorded using one electrode that was placed beneath each participant’s eye. These data were preprocessed with the Matlab AMRI EEG fMRI Toolbox v0.1.4 (99) to remove gradient and ballistocardiogram artifacts, and were then further high-pass filtered with a cut-off frequency of 0.1 Hz to remove slow drifts using the MNE-Python v1.12.1 toolbox (105). Functional MRI triggers were used to time-lock the beginning of each epoch, creating one epoch per TR (2.1s). For each epoch, the mean signal amplitude was computed, yielding one value per functional volume. The framewise difference of the resulting signal was computed and rectified (absolute value), yielding a regressor that captured how much the eyes moved between volumes, as assessed by EOG. Finally, a two-point moving average was taken to temporally align the signal with our eye voxel-based gaze predictor. The resulting EOG-based regressor was then residualized with respect to head motion in the same way the eye voxel-based regressor was (six rigid-body parameters, their temporal derivatives, and FD) and negative deflections were again clamped to 0. This EOG-based regressor was then fit to the time series of each brain voxel using the same GLM configuration as our eye voxel-based regressor. Note that 5 runs were excluded of one participant (sub-16) due to missing EEG data.

### 11) Retinotopic regions-of-interest analysis

In addition to our voxel-wise results (Figure 2), we performed a post-hoc region-of-interest (ROI) analysis for atlas-defined retinotopic regions (28). This analysis characterized the distribution of gaze-dependent brain activity across regions typically associated with vision. Participant-level *t*-statistic maps were projected to the FSAverage pial surface before extracting the mean across each ROI using Nilearn’s *SurfaceLabelsMasker*. A two-tailed sign-flipping permutation procedure across participants (10,000 permutations, one-sample *t*-test) tested the *t*-statistic in each ROI against zero for both Wake and Sleep. Derived *p*-values were corrected for multiple comparisons across ROIs using a False Discovery Rate with *q* = 0.05. The *t* statistic distributions of each ROI for both Wake and Sleep were visualized with whisker box plots (Figure 2B).

### 12) Functional connectivity analysis

To examine the relationship between gaze behavior and functional connectivity (FC) estimates in the brain, we computed FC matrices for the Schaefer 500 parcels (34) with and without partialling out the effect of inferred gaze dynamics from each brain voxel’s time series. In both analyses, the six rigid-body motion parameters, their derivatives, framewise displacement, cerebrospinal fluid, and white matter were used as nuisance regressors to denoise the voxel time series. In addition, one of the analyses included the raw and HRF-convolved gaze predictor to account for additional gaze-related variance. To maximize sensitivity and reduce noise, we combined both Wake and Sleep in this analysis, and excluded runs with fewer than 30 functional volumes in a given cognitive state (35). Pair-wise Pearson correlation was then computed for all parcels to obtain run-level FC matrices, which were then Fisher z-transformed and averaged within each participant, yielding one FC matrix per state and participant. We then compared FC matrices with and without the gaze predictor edge-wise using a sign-flipping permutation test (10,000 permutations, mean difference test statistic, sign-flipped at the participant level) against the null hypothesis of zero mean difference with a two-tailed test, correcting for multiple comparisons using the Benjamini-Hochberg False Discovery Rate at *q* = 0.05. The difference between FC matrices was then visualized alongside significant edges (Figure 4).

## Supplementary Material

**Figure S1:**
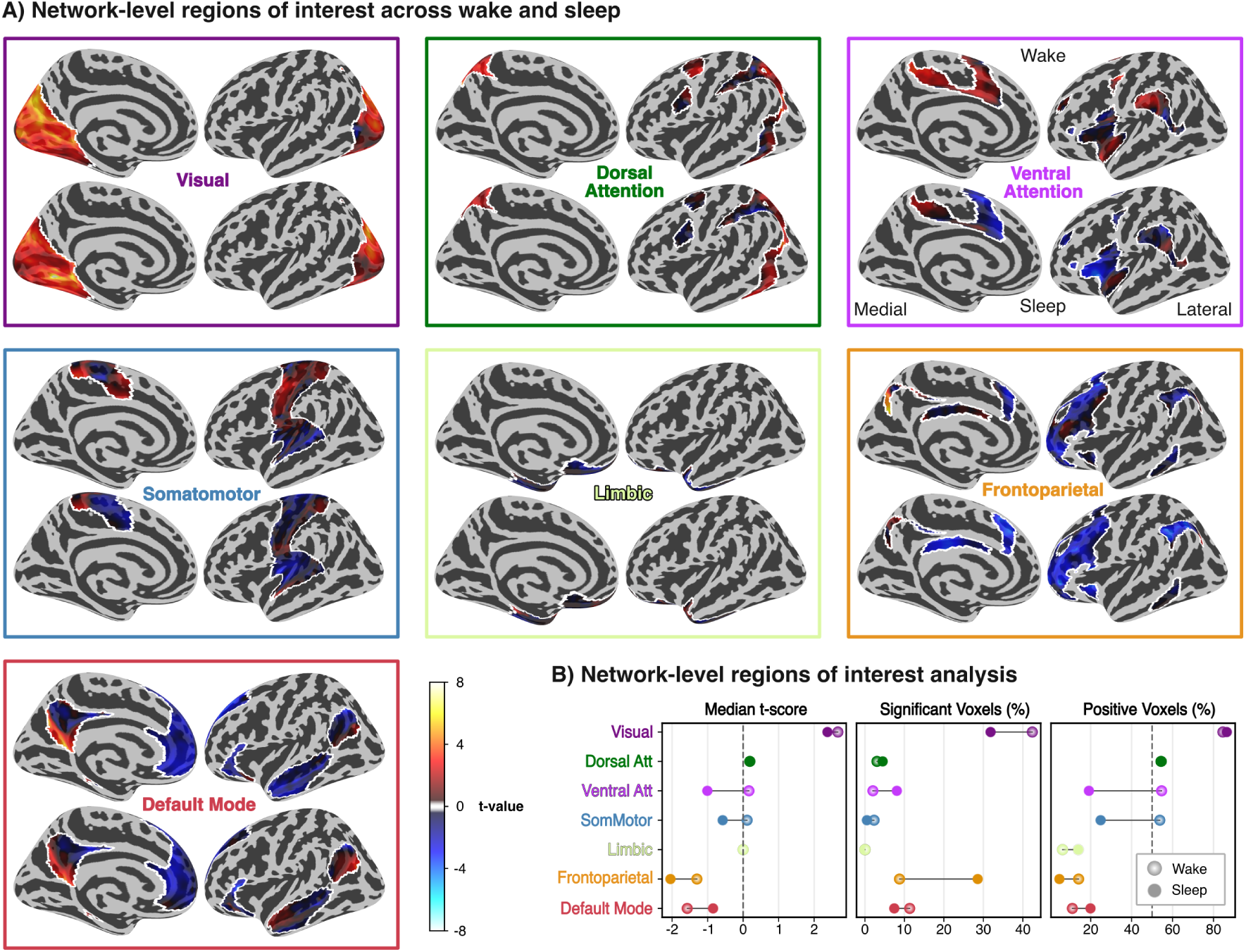
Network-level summary of gaze-dependent activity patterns for Wake vs. Sleep. (A) Network-level region-of-interest maps displaying voxel-wise general linear model results estimated for the eye voxel-based gaze predictor across Wake and Sleep. Maps display group-level results of one sample t-tests performed against zero using FSL’s FLAMEO mixed-effects model (FLAME 1+2) projected onto FreeSurfer’s FSaverage surface and masked with each of the 7 Yeo networks (36). (B) Post hoc network-level summary across states. Descriptive metrics of voxels in the group-level t-statistic maps for Wake and Sleep were computed within each of the 7 Yeo networks (36). Left panel: median t-score across voxels in each network. Center panel: percentage of voxels in significant clusters surviving False Discovery Rate correction (*q* < 0.05, cluster extent > 20 voxels). Right panel: percentage of positive t-scores, with the dotted line indicating the inflection point where positive values become the majority.

**Table S1:**
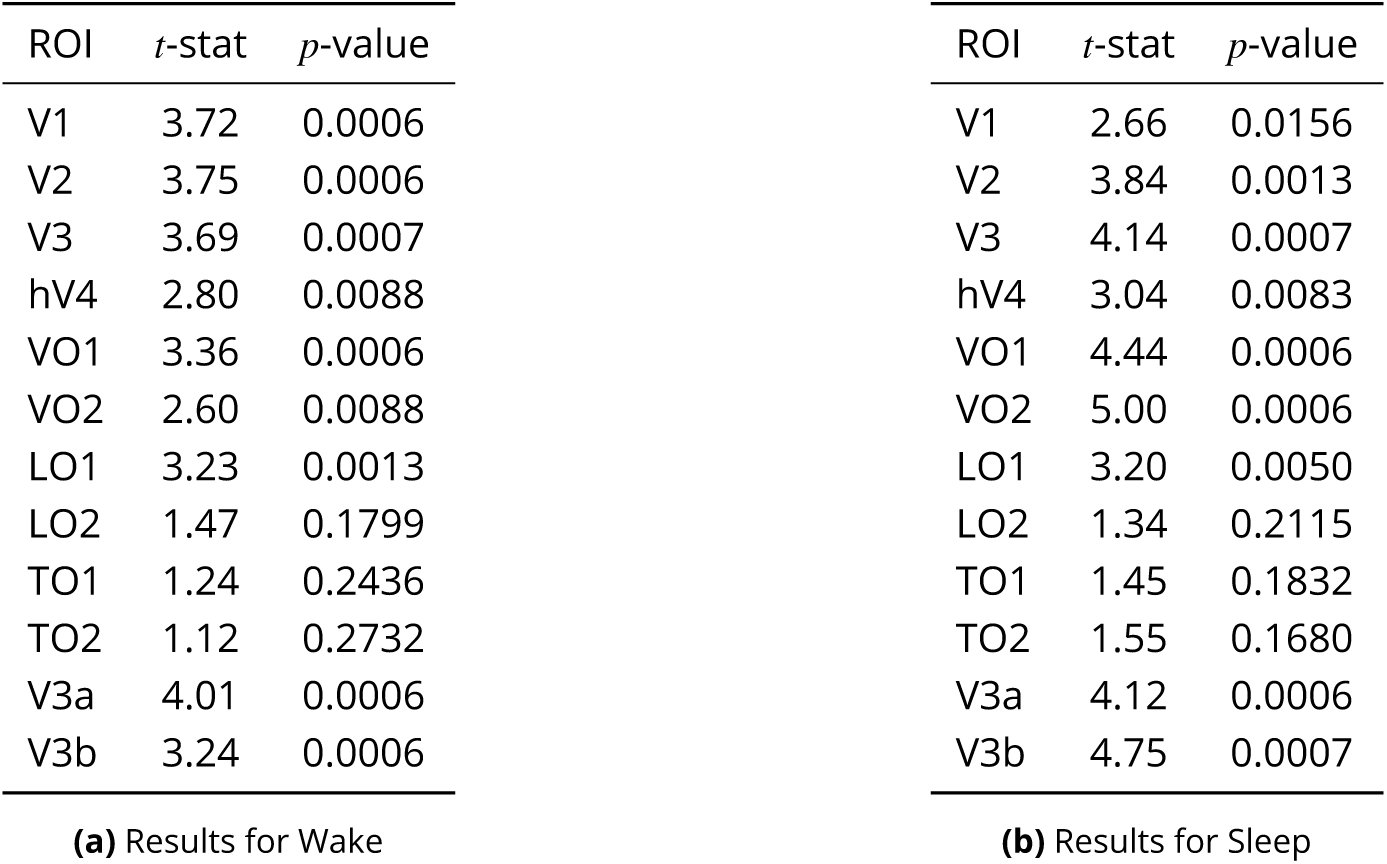
Post-hoc region-of-interest analysis for Benson atlas defined retinotopic visual areas across Wake and Sleep. (**28**). Tables display the *t*-statistic and *p*-value (sign-flipping permutation test, 10,000 permutations, one-sample *t* test, FDR corrected at *q* = 0.05) of mean participant-level *t*-statistics within each ROI. Note that these statistics are reported for the sake of completeness but do not reflect independent evidence as the analyses were performed after the voxel-wise results were known.

| ROI | *t*-stat | *p*-value | ROI | *t*-stat | *p*-value |
| --- | --- | --- | --- | --- | --- |
| V1 | 3.72 | 0.0006 | V1 | 2.66 | 0.0156 |
| V2 | 3.75 | 0.0006 | V2 | 3.84 | 0.0013 |
| V3 | 3.69 | 0.0007 | V3 | 4.14 | 0.0007 |
| hV4 | 2.80 | 0.0088 | hV4 | 3.04 | 0.0083 |
| VO1 | 3.36 | 0.0006 | VO1 | 4.44 | 0.0006 |
| VO2 | 2.60 | 0.0088 | VO2 | 5.00 | 0.0006 |
| LO1 | 3.23 | 0.0013 | LO1 | 3.20 | 0.0050 |
| LO2 | 1.47 | 0.1799 | LO2 | 1.34 | 0.2115 |
| TO1 | 1.24 | 0.2436 | TO1 | 1.45 | 0.1832 |
| TO2 | 1.12 | 0.2732 | TO2 | 1.55 | 0.1680 |
| V3a | 4.01 | 0.0006 | V3a | 4.12 | 0.0006 |
| V3b | 3.24 | 0.0006 | V3b | 4.75 | 0.0007 |

(a) Results for Wake
(b) Results for Sleep

**Figure S2:**
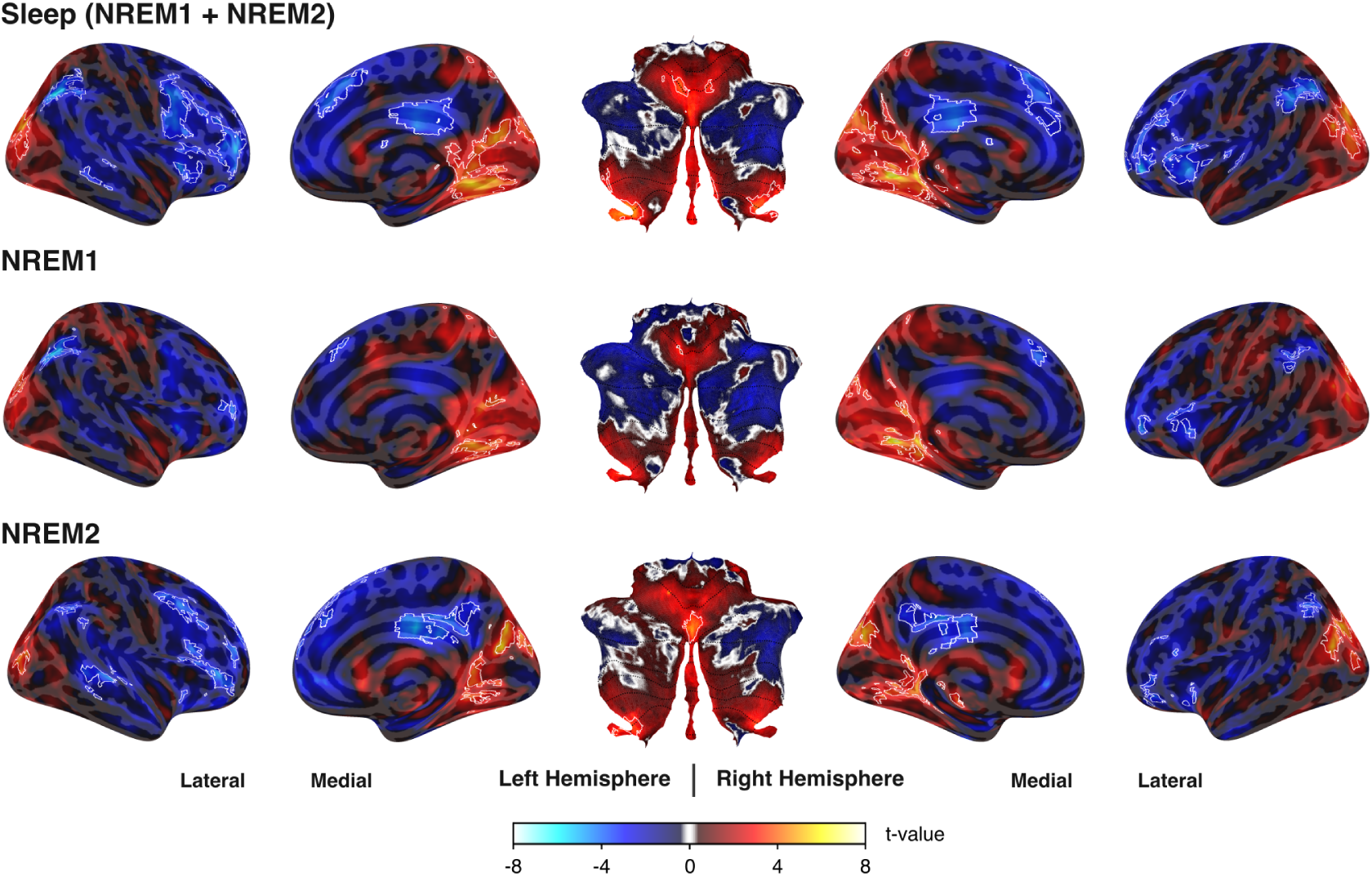
Gaze-dependent brain activity across sleep stages. Surfaces display voxel-wise general linear model estimates for the eye voxel-based gaze predictor, shown separately for Sleep (combined NREM1 and NREM2), NREM1, and NREM2. Maps show group-level results of one sample t-tests performed against zero using FSL’s FLAMEO mixed-effects model (FLAME 1+2), with cerebral results projected on FreeSurfer’s FSaverage surface and cerebellar results mapped onto SUITPy’s flatmap surface. Exploratory analyses in the subset of participants with sufficient data in NREM1 and NREM2 showed no significant differences in the mean amplitude or standard deviation of the gaze predictor between Wake and NREM1 or between Wake and NREM2 (one-sample t-tests on pseudo z-scores from within-participant state-flipping permutation tests, all *p* > 0.05, *n* = 24). Mean amplitude was nevertheless slightly lower in NREM2 than in NREM1 (group-level pseudo *z* − *score* = 0.55, *t*(23) = 2.54, *p* = 0.019), suggesting that participants performed fewer and/or smaller eye movements in deeper sleep. Additionally, NREM1 gaze-dependent brain activity displayed fewer and smaller significant clusters than both Wake and NREM2, indicating that divergence between Wake and Sleep is unlikely to originate in NREM1.

**Figure S3:**
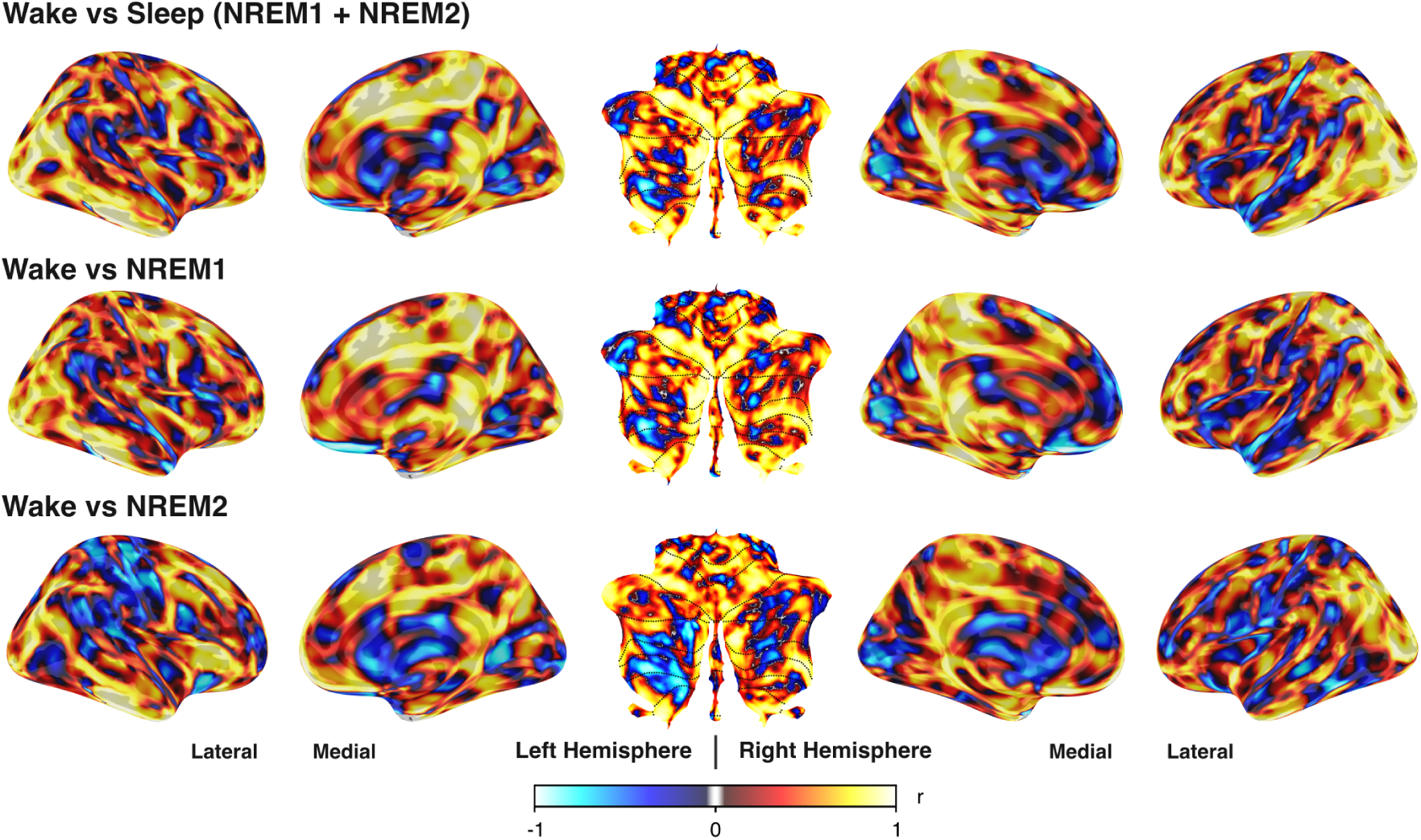
Searchlight-based comparison of gaze-dependent activity across cognitive states. Statistical maps show the local similarity of group-level t-statistics across Wake vs Sleep (combined NREM1 and NREM2), Wake vs NREM1, and Wake vs NREM2. The maps were compared using Pearson correlation (r) within local geodesic searchlights of 23*mm* radius for the cerebrum and 11*mm* for the cerebellum. Positive values (warm colors) indicate spatial consistency across states, whereas negative values (cool colors) indicate localized differences in eye–brain dynamics between states. Between state-pairs revealed that Wake and NREM1 were more similar to each other (median vertex correlation cerebrum *r* = 0.44, cerebellum *r* = 0.47) than Wake and NREM2 (median vertex correlation cerebrum *r* = 0.31, cerebellum *r* = 0.40).

**Figure S4:**
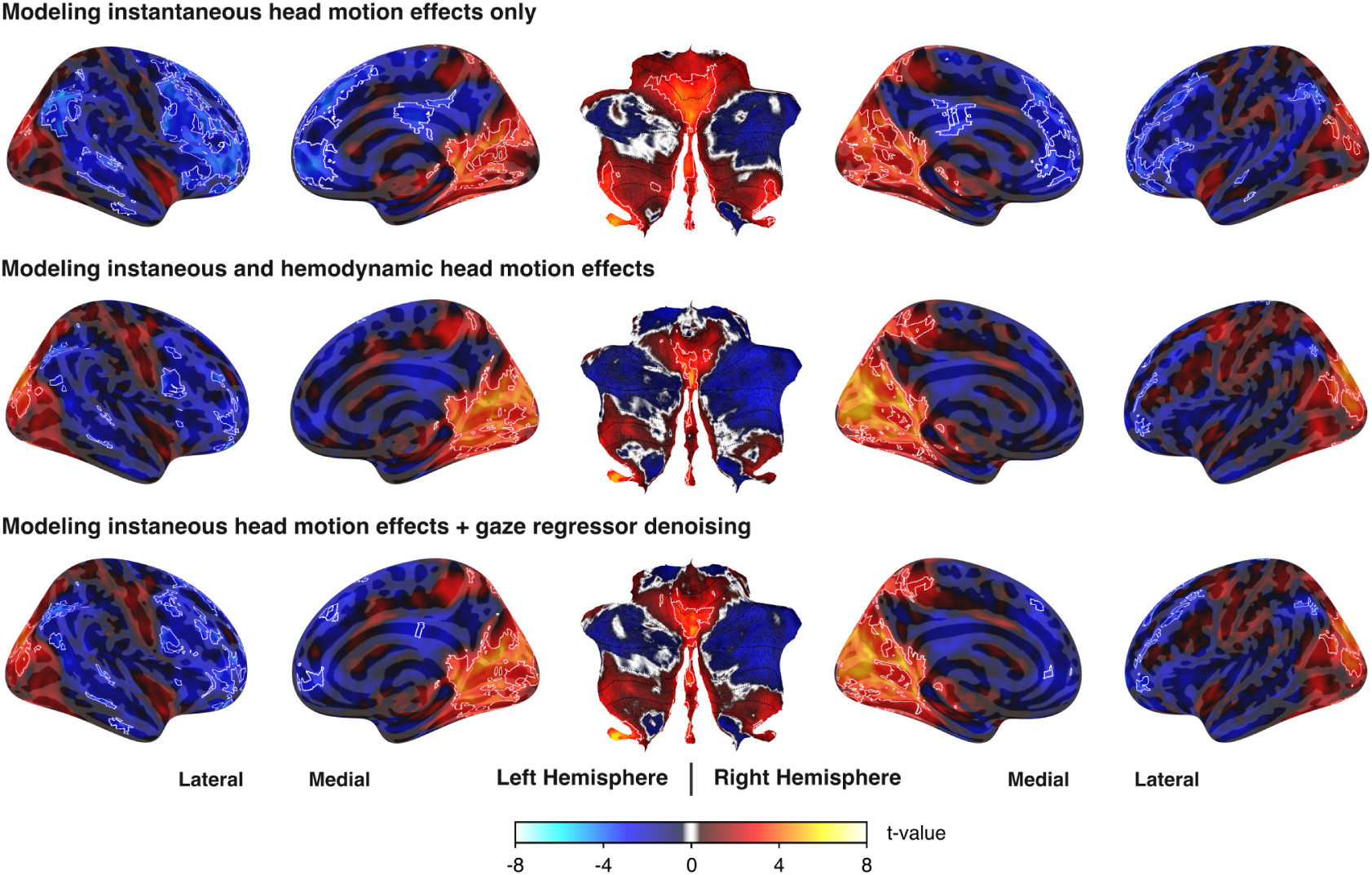
Gaze-dependent brain activity under different head motion denoising strategies. Surfaces display voxel-wise general linear model estimates for the eye voxel-based gaze predictor (combined Wake and Sleep) for different head motion denoising strategies. Maps show group-level results of one sample t-tests performed against zero using FSL’s FLAMEO mixed-effects model (FLAME 1+2), with cerebral results projected on FreeSurfer’s FSaverage surface and cerebellar results mapped onto SUITPy’s flatmap surface. Top panel: modeling only the instantaneous effects of head motion (incl. six rigid-body head motion parameters, their derivatives, and framewise displacement). Middle panel: modeling both the instantaneous and hemodynamic (framewise displacement only) effects of head motion. Bottom panel: modeling only instantaneous head motion effects but denoising the gaze predictor of head motion (six rigid-body head motion parameters, their derivatives, and framewise displacement) before including in the GLM design matrix.

**Figure S5:**
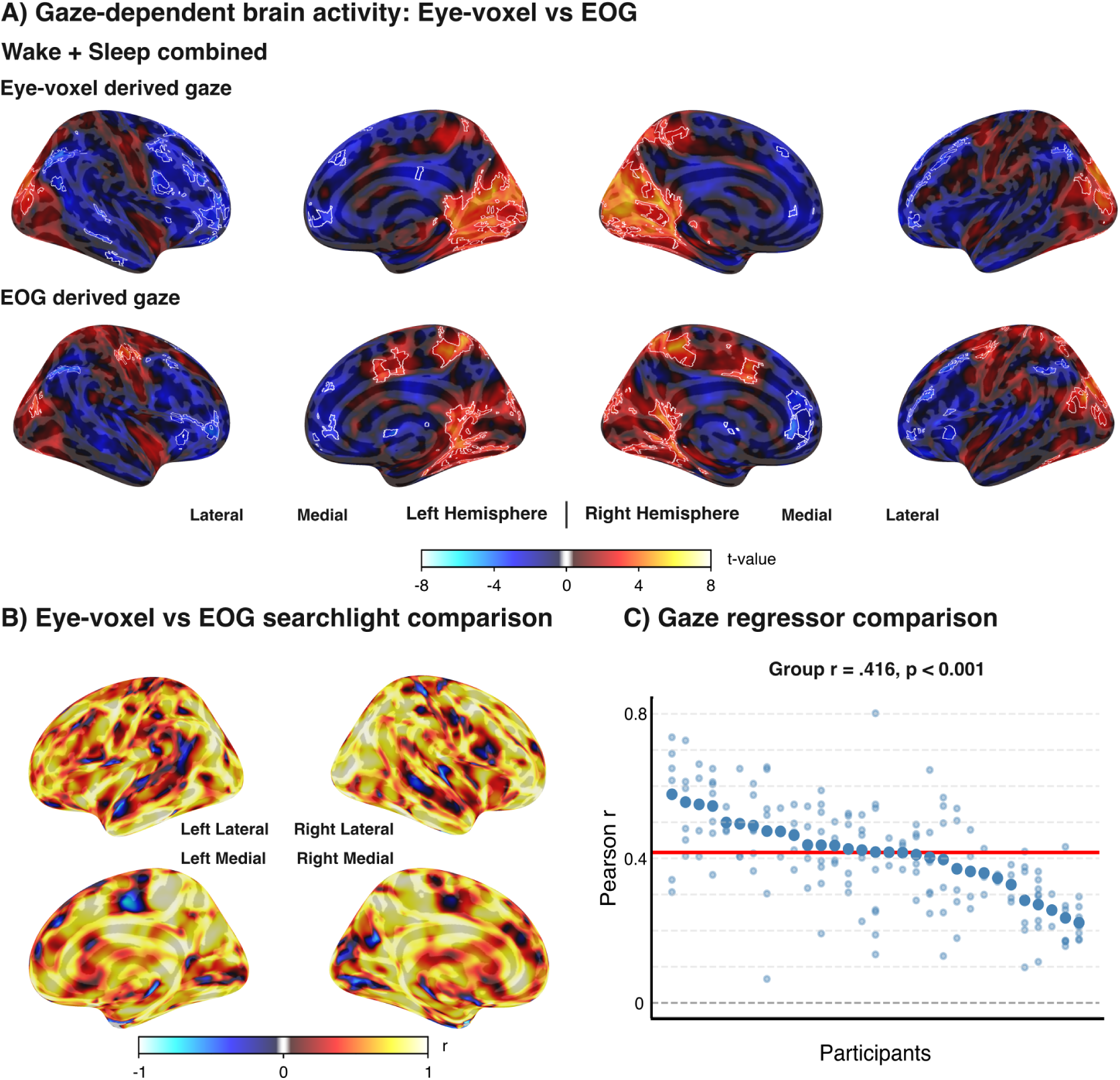
Eye voxel-based and EOG-based gaze predictor agreement. (A) Comparison of eye-voxel derived vs EOG derived gaze-dependent brain activity. Surfaces display voxel-wise general linear model estimates for the eye voxel-based gaze predictor for voxel and EOG derived gaze predictor, fit in two separate but equivalent models. Maps show group-level results of one sample t-tests performed against zero using FSL’s FLAMEO mixed-effects model (FLAME 1+2) projected on FreeSurfer’s FSaverage surface. White contours indicate clusters which survived False Discovery Rate correction (*q* < 0.05, cluster extent > 20 voxels). (B) Surface searchlight-based comparison across eye-voxel and EOG derived maps. Statistical maps show the local similarity of group-level t-statistics across eye-voxel and EOG derived results. The maps were compared using Pearson correlation (r) within local geodesic searchlights of 23*mm* radius and show a large spatial overlap (median vertex *r* = 0.63). (C) Pearson correlation between the voxel and EOG derived gaze predictor amplitudes. Large dots represent participant means (Fisher-averaged across runs) in descending order. Small dots indicate individual runs. The red horizontal line shows the group mean (*r* = 0.418, one-sample t-test: *p* < 0.001).

